# Monkeys and Humans Implement Causal Inference to Simultaneously Localize Auditory and Visual Stimuli

**DOI:** 10.1101/823385

**Authors:** Jeff T. Mohl, John M. Pearson, Jennifer M. Groh

## Abstract

The environment is sampled by multiple senses, which are woven together to produce a unified perceptual state. However, optimally unifying such signals requires assigning particular signals to the same or different underlying objects or events. Many prior studies (especially in animals) have assumed fusion of cross-modal information, whereas recent work in humans has begun to probe the appropriateness of this assumption. Here we present results from a novel behavioral task in which both monkeys and humans localized visual and auditory stimuli and reported their perceived sources through saccadic eye movements. When the locations of visual and auditory stimuli were widely separated, subjects made two saccades, while when the two stimuli were presented at the same location they made only a single saccade. Intermediate levels of separation produced mixed response patterns: a single saccade to an intermediate position on some trials or separate saccades to both locations on others. The distribution of responses was well described by a hierarchical causal inference model that accurately predicted both the explicit “same vs. different” source judgements as well as biases in localization of the source(s) under each of these conditions. The results from this task are broadly consistent with prior work in humans across a wide variety of analogous tasks, extending the study of multisensory causal inference to non-human primates and to a natural behavioral task with both a categorical assay of the number of perceived sources and a continuous report of the perceived position of the stimuli.

**Author Summary:** We experience the world through multiple sensory systems, which interact to shape perception. To do so, the brain must first determine which pieces of sensory input arise from the same source and which have nothing to do with one another. To probe how the brain accomplishes this causal inference, we developed a naturalistic paradigm that provides a behavioral report both of the number of perceived stimuli and their locations. We tested performance on this task in both humans and monkeys, and we found that both species perform causal inference in a similar manner. By providing this cross-species comparison at the behavioral level, our paradigm lays the groundwork for future experiments using neuronal recording techniques that may be impractical or impossible in human subjects.

## Introduction

Perception is inherently multisensory. Often, information from one sensory modality can reduce uncertainty about another, such as reading the lips of a speaker to improve speech comprehension [1]. However, combining such visual and auditory cues is only beneficial if they originate from the same source in the environment. In the lip-reading example above, the observer must correctly pair the sight of the lip movements with the sound of the person speaking. This is an example of a causal inference (CI) problem: determining which source(s) are most likely to have caused specific sensory observations.

Recent behavioral studies in humans have modeled such multisensory perception hierarchically, involving an assessment of the relative likelihood of two causal scenarios (same source or different sources) which then influences how sensory information is interpreted to perform a given behavioral task (e.g. localization of a particular stimulus source) [2–11]. In contrast, previous multisensory neurophysiological research in animals has generally assumed that the animals fuse the visual and auditory sources [12–15]. This discrepancy of approach has limited the inferences that can be drawn connecting neural and behavioral observations, including at the theoretical level [16–19]. One challenge has been that, to date, the tasks that have been used to probe causal inference in humans have involved behavioral reports that are arbitrarily associated to the stimulus at hand, such as using button presses or cursor movements to indicate the location of a sound and/or visual stimulus [2–4,14]. These scenarios pose challenges for animal training, where it is generally easier to shape naturally occurring responses than induce arbitrary stimulus-response associations de novo.

Accordingly, in this study we developed a task that can be deployed in both humans and animals, to provide a direct comparison of multisensory causal inference across species. We leveraged an innate behavior with an intuitive task (localizing the source of a stimulus) and response type (orienting to said source using saccadic eye movements). By requiring subjects to report both auditory and visual targets on each trial by making saccades to the perceived source of each stimulus, we could ascertain whether they perceived the stimuli to be fused vs. segregated and where exactly they perceived them to be. This provided reports of both explicit (number of saccades) and implicit (location of saccades) causal inference on each trial.

We find that monkey and human subjects behave similarly, and that their behavior reflects similar causal inference strategies. Subjects tended to make one saccade when the visual and auditory stimuli were located in the same spatial position and two saccades when they were widely separated (e.g. greater than 12 degrees apart). The transition between these modes was well described by models that incorporate causal inference, consistent with previous reports concerning human performance in other similar tasks [2–5]. These results suggest that this single/dual saccade task provides a reliable assay of multisensory causal inference that can be deployed in both humans and animal models, and sets the stage for future work bridging between the neural and behavioral domains.

## Results

### Behavioral paradigm

Subjects (7 human and 2 monkeys) were seated in a dark, anechoic chamber facing a row of co-located speakers and LEDs. Trials were randomly interleaved and consisted of either unisensory (single auditory or visual stimulus), or multisensory (both modalities, presented at the same time and for the same duration) stimuli. Multisensory trials had varying amounts of spatial separation between the auditory and visual targets, ranging from 0 degrees (coincident) to 36 degrees. On every trial, subjects were required to report the location of both the auditory and visual stimulus. For unisensory or multisensory-coincident trials (Fig 1A, top panel), subjects made a single saccade to the perceived location of the stimulus source and then held fixation at that point. For multisensory-separate trials, subjects made two saccades in rapid succession, one to each of the perceived sources. It is important to note that data were not excluded if subjects failed to make two saccades in this condition, as it is expected that subjects will often perceive the two stimuli as coincident if the spatial separation is small (6-12 degrees). This dual report design allowed characterization of both explicit (one vs. two saccades) and implicit (location of fused percept or independent percepts) causal inference on each trial.

**Fig 1.**
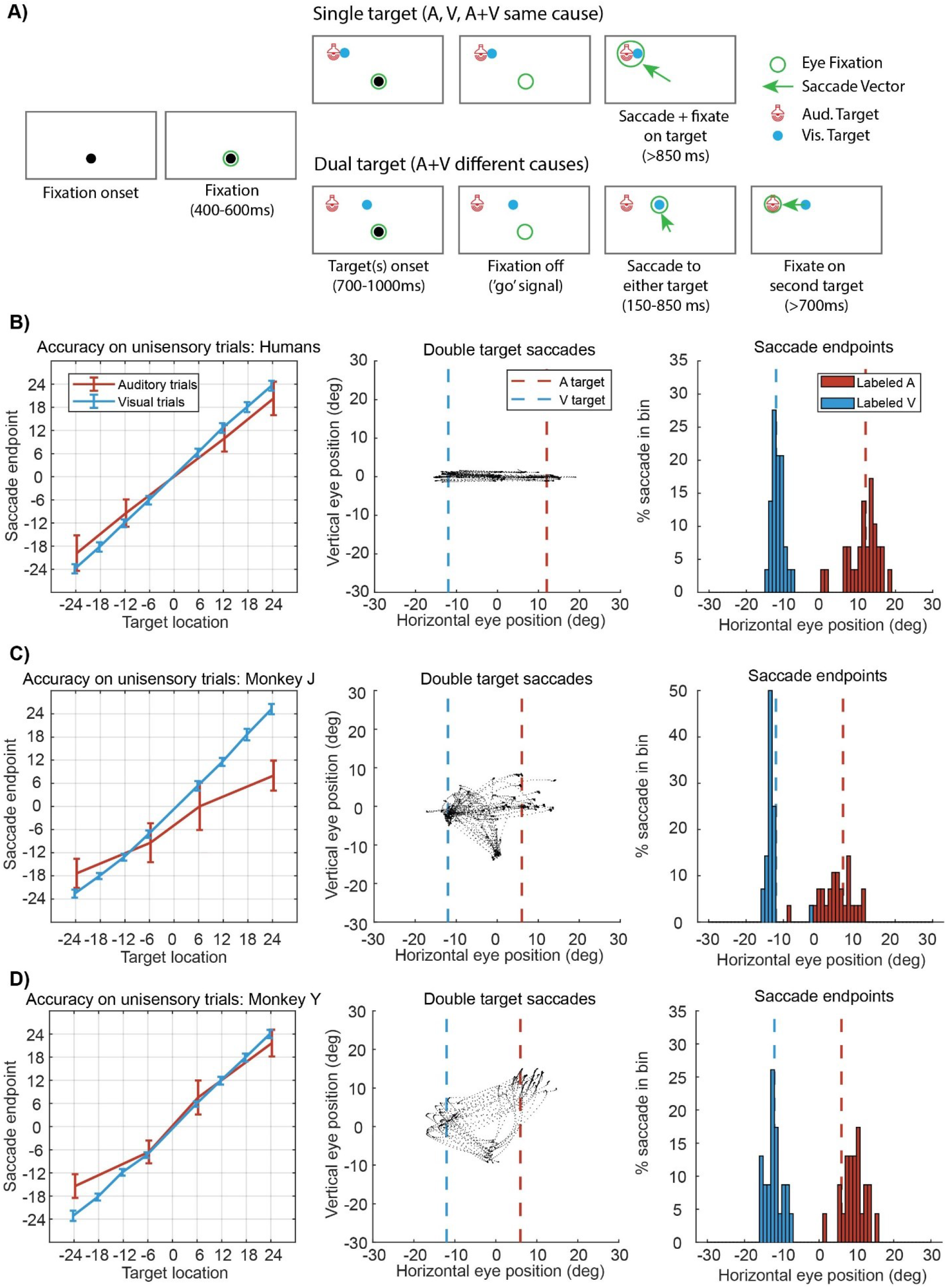
Subjects accurately localize multiple stimuli in a novel behavioral paradigm. **A)** Each trial begins with fixation at a central target location. After a variable stimulus presentation interval, subjects respond by making saccades to the sensory target(s). For single target trials (top: either unisensory trials or trials with coincident auditory and visual stimuli), subjects make a single saccade to the perceived location. For multiple target trials (bottom), subjects make two saccades in rapid succession to each target in any order. These two trial types and all target combinations were interleaved throughout the session. **B)** (Left) Human subjects were able to localize both visual (blue) and auditory (red) stimuli at all stimulus locations when presented alone on unisensory trials. Note that auditory responses have higher standard deviation, indicating lower perceptual accuracy (error bars show std. dev.). Example eye traces (center) and extracted saccade endpoints (right) for a representative dual stimulus condition with targets well separated in space (24 degrees separation). In the right panel, saccade endpoints are color coded according to whether they were labeled as auditory or visual responses on the given trial (see methods). **C)** and **D)** same as in B but for two monkey subjects. Target separation for center and right panels is 18 degrees (rather than 24 degrees for humans) due to a difference in targets used between species (see methods).

First, we sought to determine whether both monkeys and humans perform the above task in a qualitatively similar manner. Subjects were able to localize both the auditory (red) and visual (blue) stimuli when presented on unisensory trials (Fig 1 B-D, left panels). Visual localization was much more accurate and less biased (across subjects SD = 1.08 ± 0.07 deg., mean absolute error = 0.49 ± 0.19 deg.) than auditory localization (SD = 4.07 ± 0.36 deg., error = 4.80 ± 1.61 deg.), consistent with the higher sensory reliability of visual information for spatial localization tasks [20,21]. On trials where the targets were well separated, subjects accurately localized both targets by making two saccades in rapid succession (Fig 1 B-D, center panels). The distribution of endpoints from these saccades were extracted and formed the basis for comparison with model predicted distributions (Fig 1 B-D, right panels). These results demonstrate that both human and monkey subjects grasped the primary goals of the task and accurately reported both visual and auditory targets on a single trial.

### Audio-Visual Causal inference

Causal inference, as it relates to multisensory localization, has two primary perceptual consequences. Explicit causal inference is the most straightforward and amounts to simply determining which of two (or more) causal scenarios is most likely to have generated the perceived sensory inputs (Fig 2A) [14,22]. The second consequence occurs when subjects localize stimuli in a scenario where the causal structure is uncertain. Each of the potential causal structures should result in different source estimates [2,10,11]. This means that any reported location is implicitly shaped by the explicit judgement, as subjects will report intermediate locations (Fig 2B, purple curves) only when they judge the stimuli to share a source, and not when they perceive them as separated (Fig 2B, red and blue curves). We designed our paradigm to provide readouts of both the explicit and implicit components on each trial.

**Fig 2.**
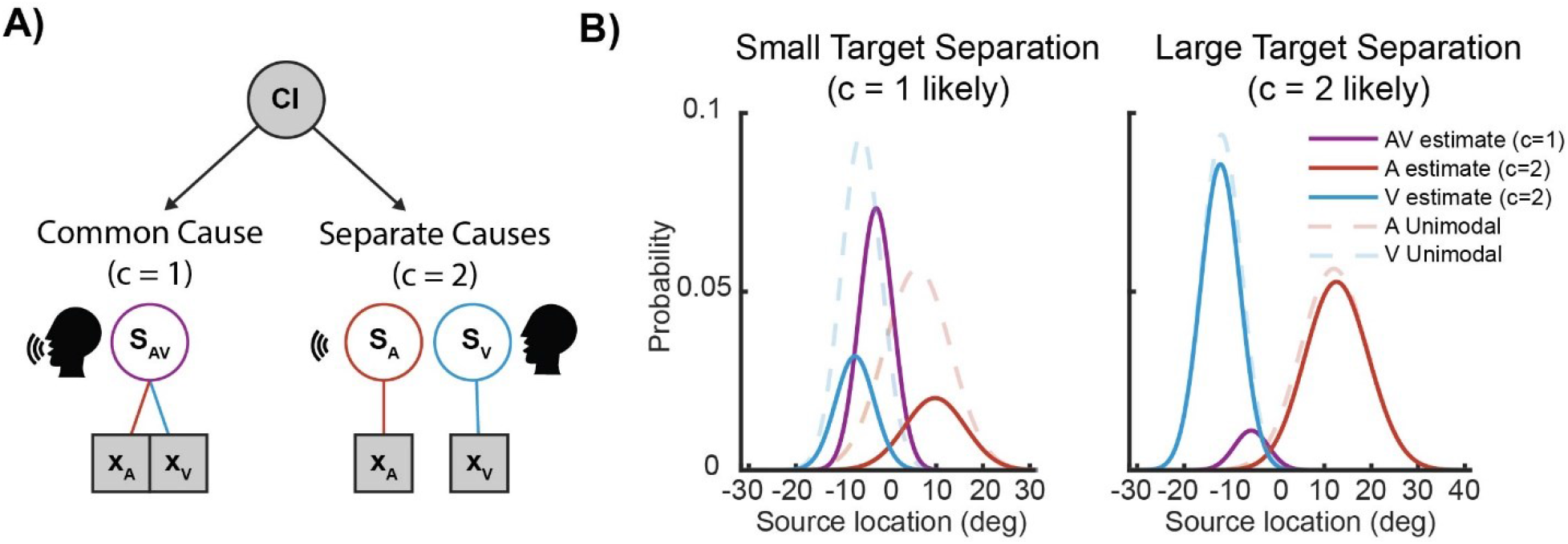
Causal inference in Audio-visual localization. **A)** The generative model producing sensory percepts (x_A_, x_V_) is assumed to have two potential causal structures: either sharing a common source (though perturbed by different amounts of sensory noise, left branch) or having independent sources (right branch). Causal inference is accomplished by inverting this generative model and determining which branch is most likely given the observations x_A_ and x_V_. **B)** Estimated probability distributions for the source of sensory percepts under the one cause (*Ŝ_AV_*, purple) or two cause (*Ŝ_A_*, red or *Ŝ_V_*, blue) conditions. The distributions are normalized such that the total area under the solid curves sum to one, so that the relative ratio between one and two saccade responses can be seen by comparing the left and right panels.

#### Modeling

We quantitatively modeled responses in our task by adapting generative models common in the literature [2,3,5]. We built a set of several observer models, which reflect different assumptions about the observer’s causal inference strategy, and compared them to a null model involving a default strategy instead. Our overarching goal was to ascertain whether and how subjects compared information across modalities to infer the number of causes, and whether they then used that information to inform their localization of those causes.

On each trial, we assumed that some set of sensory sources, *S_A_* and *S_V_*, produced noisy internal measurements *x_A_* and *x_V_*. These noise distributions were assumed to be Gaussian, centered on the source, with sensory variance encoded by the free parameters 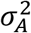 and 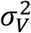, respectively. We assumed that these sensory variances were used for both the generative model of responses and the observer’s internal estimate of causality, that is, both the localization and source judgement steps. Both auditory and visual stimuli were assumed to share the same, normally distributed prior, centered at 0 with variance 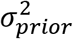.

We considered two separate response strategies for performing the unity judgement component of the task based on these noisy internal measurements: do subjects use the visual and auditory information on each trial or do they use a default heuristic that amounts to guessing in a probabilistic fashion? The first possibility is captured under an idealized Bayesian observer (Bay, see methods). For the Bay model, the observer reports a common cause when the posterior probability for that one cause is greater than 0.5, Pr(*c* = 1|*x_A_*, *x_V_*) > 0.5. The prior probability of common cause, *p_common_* = Pr(*c* = 1) is a free parameter.

The default, heuristic possibility is captured under a non-causal-inference probabilistic fusion model (PF). This model does not take the disparity between the visual and auditory stimuli into consideration at all. Rather, participants might always fuse, always segregate, or make a probabilistic choice between the two (the latter indicating that the subject has learned that a mix of behaviors is required). The PF model is equivalent to forced fusion or forced segregation models commonly compared with causal inference models [2,3,23], but is more general with the possibility of a response pattern intermediate to these extremes. For the PF model the observer reports a single cause on some fixed percentage of the trials, defined by *P_common_*.

Estimating the source location, *p*(*Ŝ* |*x_A_*,*x_V_*), depends on the assumed causal structure. If the measurements *x_A_* and *x_v_* are assumed to originate from the same source, the localization estimate will be a weighted average reflecting the relative reliability of each of the cues (*Ŝ_AV_*, Fig 2B, purple), consistent with well established maximum likelihood models of multisensory integration [20,24]. When the sources are assumed to be independent, the resulting estimates are independent and rely only on the sensory information and the prior (*Ŝ_A_*, *Ŝ_V_*, Fig 2B, blue and red). Because the trials were interleaved with varying amounts of target separation, subjects could not know ahead of time which response pattern was ideal. This forced them to adopt some kind of behavioral strategy to arbitrate between these two response patterns (Fig 2B, left vs. right panels).

In order to characterize the localization component of the task under different behavioral strategies, we considered three possible models, two that incorporate causal inference and one that does not. The first causal inference model was a Bayes optimal strategy, in which the observer combined the potential localization estimates (i.e. fused or separate) according to the posterior probability of causal structure, Pr(*c* = 1|*x_A_*, *x_V_*), which we refer to as model averaging (MA) [2]. The second causal inference model, which we refer to as model selection (MS), implemented a heuristic decision rule [4]. Instead of weighting the fused and separate estimates according to posterior probability, the model implemented a threshold decision and simply selected whichever causal structure was most likely (i.e. selecting the fusion strategy when Pr(*c* = 1|*x_A_*, *x_V_*) > 0.5, and otherwise using the segregated strategy). For comparison with these two causal inference variants, we again compared with a default probabilistic fusion (PF) model, which does not incorporate information from the causal judgement. Instead, the subject is assumed to choose between fused and segregated response distributions randomly at some fixed rate, defined by *p_common_*. This includes the possibility for either always-fuse or always-segregate response patterns. Importantly, all of these models implement the same rule for choosing to make either one or two saccades on a given trial (the Bay model for unity judgement). This means that each model was compared only based on how well it captured the distribution of location reports, rather than the ratio of single to dual saccade reports.

We fit the models using a maximum-likelihood strategy, estimating the parameters that provided the best fit the behavioral data (number of saccades or location of saccades, for unity judgement and localization models respectively) for each condition. In addition to fitting to each of these separate task components independently, we also fit models jointly to both components of the task (i.e. maximizing likelihood for both the number of saccades and the location of saccades). When fitting the joint models, the unity judgement component was assumed to follow the Bay strategy, while the localization component was varied between the three possibilities described above. For illustration purposes only the jointly fit Bay-MA model is shown in figures 3 and 4, though all models are compared quantitatively in figure 5.

**Fig 3.**
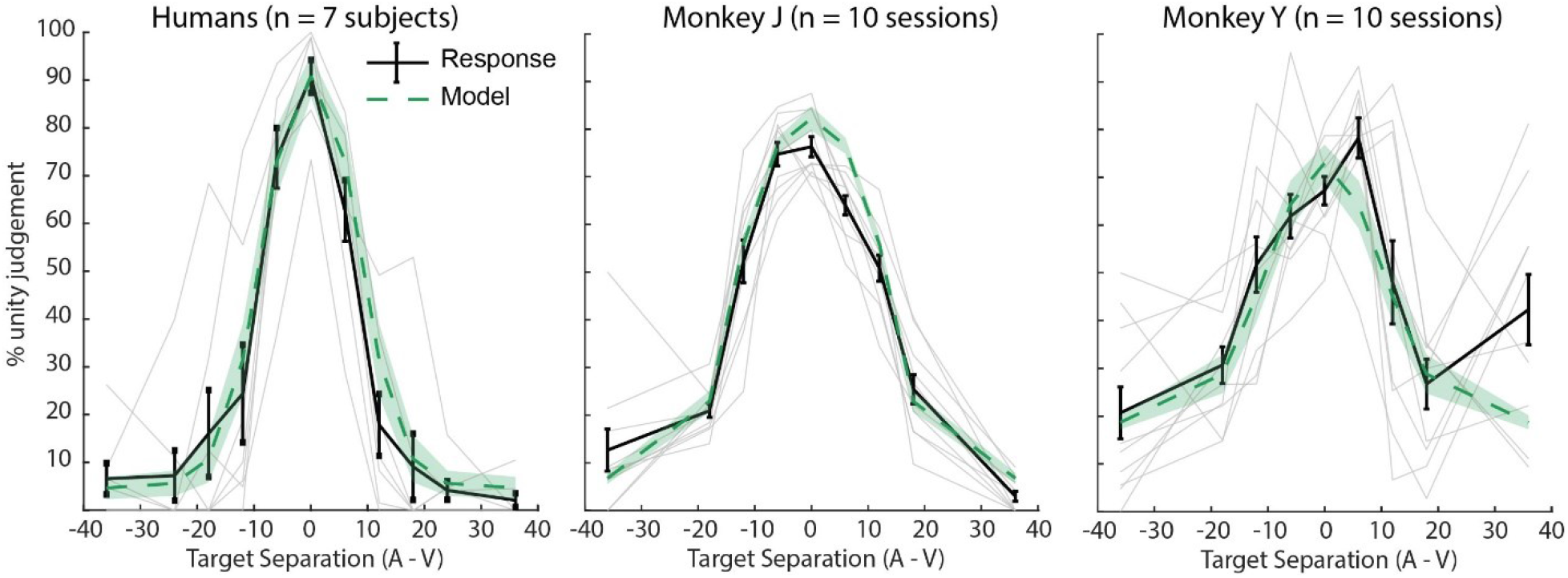
Unity judgement as a function of target disparity. Human subjects (n=7, left) and monkey subjects (n=10 experimental sessions, center, right) demonstrated a pronounced preference for making one saccade when targets were close together, rather than well separated. These judgements were well fit by a Bayesian model of causal inference (Bay-MA, green curves). Grey lines show mean responses for individual subjects (humans) or experimental sessions (monkeys). Error bars and shaded region represent SEM across subjects/sessions.

**Fig 4.**
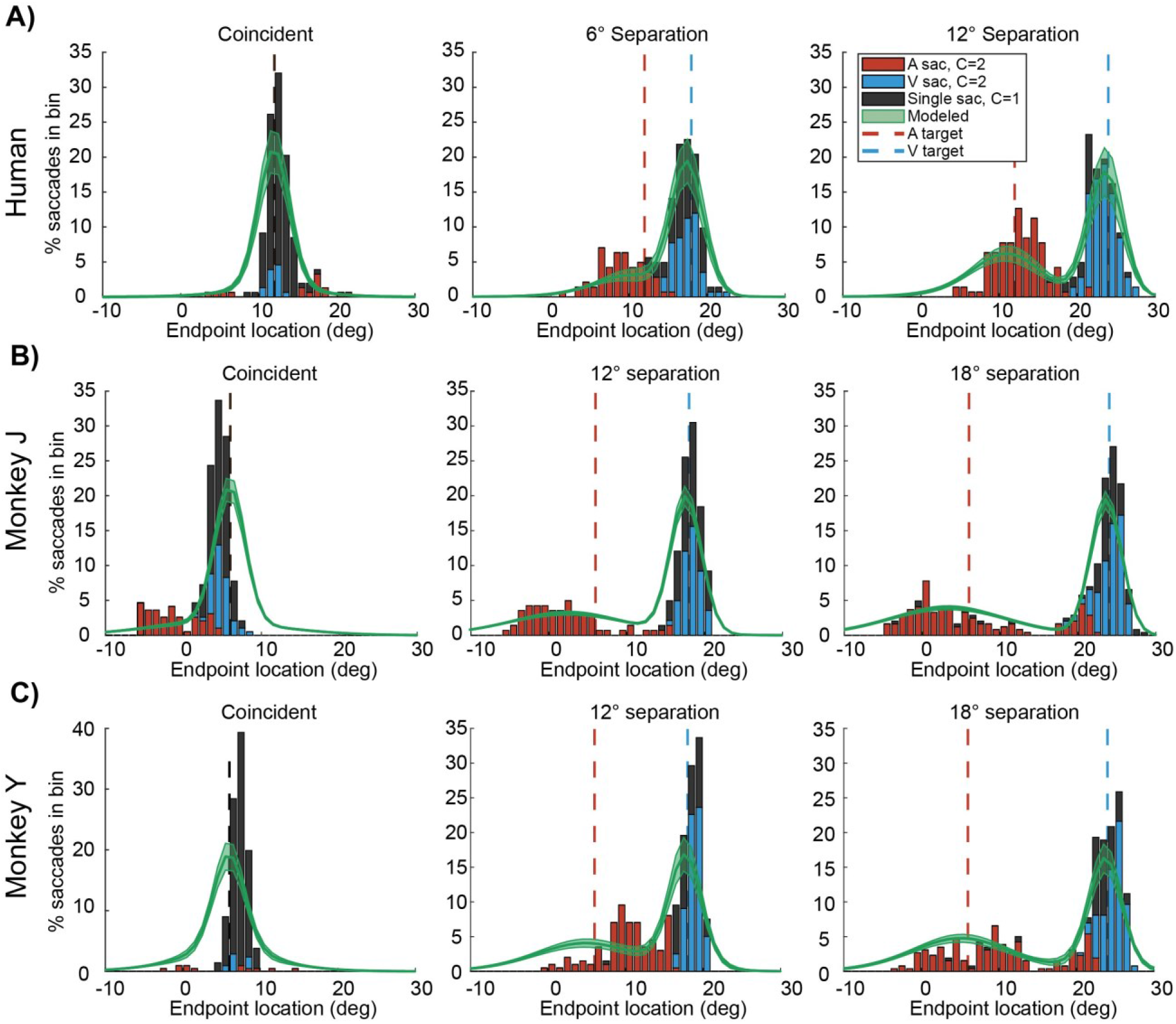
Localization of stimulus sources. Reported location is shown combined across single and dual saccade trials, with single saccade trials labeled in black while dual saccade trials contribute to both auditory (red) and visual (blue) distributions. At coincident target locations, subjects usually reported with a single saccade at that location (left panels, black bars). As the targets move further apart in space, subjects gradually shift from an integration strategy to complete segregation (center and left panels). Humans and monkeys had similar behavioral performance, though monkeys had a more pronounced auditory bias and worse localization accuracy (consistent with differences seen in unisensory trials, Fig 1 C-D). Shaded region reflects SEM.

**Fig 5.**
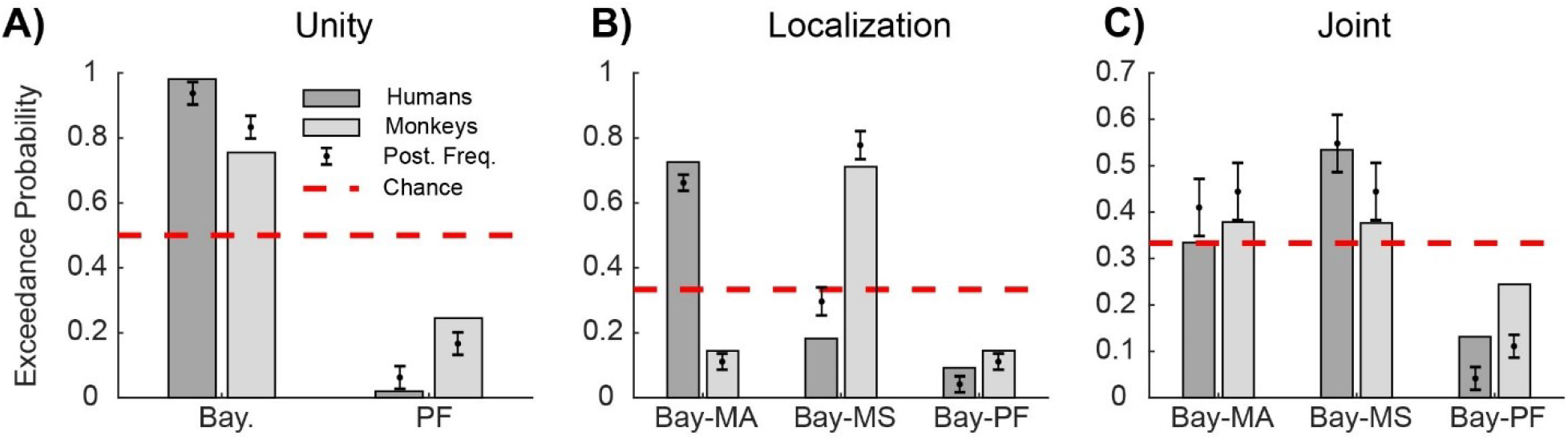
Model comparison. Protected exceedance probabilities (bars) and expected posterior model frequencies (mean ±SD) are shown for the two unity judgement and three localization models across species. **A)** Baysian CI (Bay) model provides better fits of subjects responses on the unity judgement component than a probabilistic fusion (PF) model. **B)** Performance of two CI models (MA and MS) compared with non-CI PF model, applied only to localization data. Both species are better fit by models incorporating CI (MA and MS models) than the non-CI model (PF), though the preferred model is different between species. **C)** Same as **B**, but now comparing models fit jointly to unity judgement and localization responses. There is little evidence favoring either CI model over the other, though both outperform the non-CI PF model.

### Unity Judgement

In order to determine whether subjects were performing causal inference in our task, we first analyzed the explicit portion of the response: whether the subject made one or two saccades. We found that subjects were much more likely to make one saccade when the targets were coincident or close together and much more likely to make two saccades when the targets were well separated (Fig 3). This means that the observers were not performing pure fusion (always integrating stimuli), nor pure segregation (always treating the stimuli as independent), but instead adopting a strategy that depended on target separation. Importantly, humans and monkeys showed qualitatively similar performance on this component of the task (Fig 3, left panel vs. central and right panels). This response pattern was well described by an ideal Bayesian observer model of causal inference for all subjects (Fig 3, Bay-MA model, green) [2]. These results indicate that monkeys understood and perform the explicit causal inference component of the task, and that their behavior is well described by a causal inference model previously only applied to human behavior.

### Localization

We next sought to determine the effects of causal inference on the localization of stimuli. When targets were presented at a single location, subjects overwhelmingly made a single saccade to that location (Fig 4, left panels, black bars). Conversely, when the targets were well separated, subjects accurately reported the location of both the visual and auditory sources (Fig 4, right panels, red and blue bars). At intermediate locations, subjects responded with a mixture of fused single saccade trials and separated double saccade trials (Fig 4, central panels). Like in the unity judgment component of the task, these response distributions were well fit by an ideal observer model performing Bayesian causal inference with model averaging (solid green line). These results demonstrate that our task recapitulates both the explicit (ratio between single and double saccade trials) and implicit (biases in localization between single and dual saccade trials) components of causal inference in both humans and monkeys.

### Model Comparison

Finally, we performed a quantitative model comparison between potential behavioral strategies for each component of the task (unity judgement and localization), as well as for models fit jointly to both components. We performed a Bayesian random-effects model comparison to determine which of the tested models provided the best fit for the observed data. In Fig 5, we report the protected exceedance probability (P_exc_, i.e. the probability that a given model is more likely than any other, corrected for chance) as well as the posterior model frequency for reference [25,26]. First, we compared the Bay model, which implements Baysian causal inference, with the null PF model in the unity judgement case (Fig 5, left). We found that the Bay model provided much better fits for both human (P_exc_ = 0.98) and monkey (P_exc_ = 0.76) subjects. Put another way, this indicates that a Bayesian causal inference strategy was ~49 times and ~3 times more likely to be the most representative model of behavior compared to a probabilistic fusion strategy for humans and monkeys respectively. These results again confirm that both species were taking cue disparity into account when reporting common or separate causes, rather than simply responding according to some fixed guessing strategy.

We compared the two different CI strategies for localization (MA and MS) as well as the non-CI probabilistic PF strategy (Fig 5, center). Both species were much better fit by one of the CI models, though the preferred strategy differed between species. The model-averaging observer provided the best fit for human subjects (P_exc_ = 0.73), while providing worse fits for monkeys (P_exc_ = 0.14). Conversely, monkeys were better fit by a model selection strategy (P_exc_ = 0.71) than humans (P_exc_ = 0.18). For both species, the probabilistic fusion model provided significantly worse fits (Human subjects: P_exc_ = 0.09; monkey subjects: P_exc_ = 0.15) than the combination of CI strategies. Collapsing across CI models, some form of CI was ~10 times more likely than the PF model for humans, while for monkeys CI was ~6 times more likely. These results indicate that subjects from both species are incorporating causal inference into the localization component of the task, rather than only the unity judgement component.

When this same set of models were fit jointly to both the localization and unity judgement data, the species level difference between model fits disappeared and it was no longer possible to differentiate the two CI strategies (Fig 5, right). Both of the CI strategies provided a better fit than the PF strategy. The PF strategy was ~7 times and ~3 times less likely than the combination of MA and MS models for humans and monkeys respectively (Human subjects: P_exc_ = 0.13; monkey subjects: P_exc_ = 0.24). This again provides evidence that both species were performing causal inference in this task, both for the localization and unity judgement components, though the exact strategy used remains an open question.

## Discussion

We have presented a novel behavioral paradigm for multisensory localization that provides rich perceptual readouts for both human and monkey subjects. We demonstrated that subjects are capable of localizing both auditory and visual stimuli on single trials where the sources of those stimuli are well separated in space. On trials where the stimuli were either coincident or separated by small amounts (6 to 12 degrees) subjects often colocalized the stimuli, reporting them as sharing the same source, consistent with previous reports of audio-visual fusion or visual capture [14,20,27–29]. Importantly, subjects shifted from fusion to segregation as a function of target separation, as predicted by existing models of causal inference [2–5]. Both human and monkey subject behavior was much better fit by models which incorporated some form of causal inference, as compared to models which spanned the possibilities from forced fusion to segregation (i.e. processing is completely separated by modality). Together these results demonstrate the effectiveness of this paradigm for eliciting causal inference judgements in both humans and monkeys and validate non-human primates as a model organism for studying the neural basis of causal inference.

These results are consistent with a growing body of behavioral research in humans indicating that causal inference is a critical component of sensory processing [2,4,5,10,11,22,30,31]. We found that relatively simple models of causal inference (with 5 or fewer parameters) provided good fits even in a complex behavioral paradigm, which had both a dual report structure and a continuous reporting variable. This further supports the relevance of such models for the study of multisensory perception, as a small number of biologically relevant parameters offer significant predictive power.

Recent human neuroimaging studies have suggested that causal inference may be accomplished by subdividing the task into pieces (i.e., integration, segregation, etc.) and performing these computations in separate brain regions before combining them in some higher level brain region such as pre-frontal cortex [11,23,32–36]. This view of hierarchical neural processing is pleasingly consistent with the hierarchical nature of ideal observer models of causal inference (Fig 2A). However, it is inconsistent with other research that showing significant interaction between modalities even in primary sensory areas [37–40], as well as numerous descriptions of multisensory integration in subcortical brain regions [12,41–47]. It is possible that this conflict is due in part to the level of experimentation, as the former findings rely on human neuroimaging (fMRI and MEG) which is necessarily limited to large-scale changes in neural activity in cortical structures, while the latter studies principally investigated single neurons or small networks of neurons. Further experiments are needed to determine whether multisensory causal inference is truly a brain-wide computation or whether it can be accomplished in smaller networks of individual neurons.

We designed this task to be compatible with monkey single unit electrophysiology experiments, which imposes some limitations from a behavioral modeling perspective. Most importantly, in order to keep the total number of conditions reasonably small, we did not vary the sensory reliability for either stimulus type. This prohibits us from weighing in on the exact nature of the causal inference, whether Bayes optimal (i.e., model averaging, which minimizes localization error) or some heuristic alternative (i.e. probability matching or comparing to a fixed criterion, which may be simpler computationally). Recent work with a more thorough model comparison has shed some light on this subject, suggesting that a heuristic fixed-criterion model may provide better fits for human behavior [5]. However, there is significant variability even between individual human subjects performing the same task [3,48], and so we leave the question of which exact model of causal inference best describes behavior to future studies which can bring more statistical power to bear.

Additionally, while using saccades as a continuous behavioral readout offers many advantages, this choice does impose certain important limitations. The first of these is that motor noise is conflated with sensory noise: that is, a portion of the variability in responses is due to errors in the motor output rather than uncertainty about the target location. Previous work has determined that the contribution of motor noise to saccade variance is only slightly smaller than the contribution of sensory noise [49], suggesting that this issue is not trivial. Therefore, our estimates of sensory noise may be higher than the actual estimates used by the subject’s brain when performing a causal inference, particularly for the visual targets which are likely to have low sensory noise. This appears to be the case when comparing response distributions to those predicted by the model (Fig 4, over dispersion of model distributions relative to histograms). However, because this is an issue that affects all compared models equally, it should not affect our model comparison results nor our conclusions.

The second limitation imposed by saccadic reporting is that there is a natural lower bound on target separation that can be realistically indicated using two saccades. Both humans and monkeys make near-constant corrective micro-saccades (<1-2 degrees in amplitude), which are necessarily differentiated from voluntary saccades in this task [50]. This makes it essentially impossible for subjects to indicate two separate targets if they perceive them as separate but closer than approximately 3 degrees apart, as any smaller magnitude of saccade would be interpreted as corrective. This is not expected to influence our results, as all target pairs used in this experiment have a separation of at least 6 degrees (or are coincident), and at this smallest separation value subjects still overwhelmingly report a single percept. However, it is worth considering in similar paradigms which might use sensory stimuli with less inherent sensory noise.

Aside from these limitations, our task has many advantages over similar, previously reported CI tasks. Localizing and orienting to sensory stimuli through saccadic eye movements is an innate behavior for primates, which facilitates comparison across species. Using saccades also allows for a continuous report of perceptual location, rather than button presses or two alternative forced choice paradigms which necessarily limit the number of potential responses and provide an indirect mapping between perception and response. Our approach therefore allows for a more thorough characterization of behavior from trial to trial, while leveraging a natural behavior to speed training and data collection compared to other continuous report tasks [14].

Another advantage is that the dual report design (requiring both a unity judgement and localization) and interleaved nature of the task is more consistent with how sensory inference is performed in the natural environment. Subjects must attend to and act on sensory information of different modalities from moment to moment. They cannot simply rely on a strategy such as focusing only on visual or auditory inputs and neglecting all others (as with a blocked trial design). This is critical for understanding the neural basis of this operation, as focusing attention on only one modality or region of space is likely to significantly influence neural responses [51].

Most importantly, this task design allows for direct comparison between human and monkey behavioral subjects, and can characterize important features of causal inference within a single experimental session. By demonstrating that monkeys and humans perform the task similarly, we validate an animal model aligned with the growing body of human behavioral research in this area. This will enable the use of much higher resolution recording techniques that are difficult or impossible to use in human subjects, which is critical for bridging the gap between our understanding of multisensory causal inference at the behavioral and neuronal levels.

## Methods

### General procedures

Human subjects (n = 7, 4 females, aged 15–30 y) were involved in this study. All procedures involving human subjects were approved by the Duke University Institutional Review Board (IRB Protocol Number: 1885). Subjects had apparently normal hearing and normal or corrected vision. Informed consent was obtained from all participants before testing, and all subjects received monetary compensation for participation.

All procedures conformed to the guidelines of the National Institutes of Health (NIH Pub. No. 86–23, Revised 1985) and were approved by the Institutional Animal Care and Use Committee of Duke University (Protocol Registry Number A115-15-04). Two adult rhesus monkeys (Macaca mulatta) participated (monkey J, and monkey Y, both female). Under general anesthesia and in sterile surgery we implanted a head post holder to restrain the head and a scleral search coil to track eye movements [52]. After recovery with suitable analgesics and veterinary care, we trained the monkeys in the experimental task.

### Behavioral paradigm

We created a novel multisensory task closely related to tasks commonly used in the literature [2–4,14]. This paradigm used a dual report design, where subjects reported both a causal judgement (one or two targets, explicit causal inference) and the target locations (implicit causal inference) on every trial.

Subjects were seated in an anechoic chamber at a distance of 1.25 m from a row of speakers and LEDs located on the horizontal plane. Eye movements were monitored via magnetic eye coil (monkeys; Riverbend) or video eye tracker (monkeys, humans; SR Research Eyelink 1000). While fixating at a central point (0 degrees horizontal, variable vertical offset of −12 to +6 degrees for monkey subjects), subjects were presented with either a light (green LED), sound (white noise), or both at one of 8 visual (± 6-24 degrees in 6 degree increments) or 4 auditory (±6 and ±24 degrees for monkeys; ±12 and ±24 degrees for humans) locations. Targets were paired such that each combination of ipsilateral pairs was used (8 pairs per side, for 16 pairs), plus 4 contralateral pairs (±12 degrees visual paired with either contralateral auditory location) for a total of 20 dual conditions. After a brief delay (600-900 ms) the fixation light was extinguished, and subjects reported percepts by making saccades to the perceived stimulus location and then maintaining fixation at that target location. On conditions with multiple targets, subjects were required to make sequential saccades to each target in any order. The timing of the task was such that subjects need to make both saccades in rapid succession, and so could not adopt a strategy of waiting until the reward was delivered (or not) before making a decision about the second saccade.

### Trial filtration and saccade detection

Trials were included as long as the subject held fixation through the go cue, and then made at least one saccade, without enforcing any restrictions on saccade accuracy. For multi-stimulus trials, trials that ended less than 600 ms after the go cue (the minimum duration for a successfully completed trial) were also excluded. This was done to minimize the number of trials that ended before the subject’s full response could be reported, and ensure that single saccades were indicative of a unified percept rather than a lapse.

Saccades were defined as any eye movement exceeding 50 degrees per second and followed by at least 30 ms of very little eye movement (max velocity <25 deg/s). Saccades of less than 3 degrees were considered corrective and were not included as responses in subsequent analyses.

### Behavioral modeling

We implemented a class of causal inference models which is common in human behavioral multisensory research (see [9] for review). These models arbitrate between two sensory processing strategies. The first strategy treats sensory stimuli as completely independent, amounting to unisensory estimation of the parameter of interest (in this case, location of the source) for each stimulus. The second implements the established maximum-likelihood form of cue integration, which has been shown to provide excellent descriptions of human behavior in conditions where the disparity between multisensory cues is small or the cues are mandatorily fused [20,24,53]. Different models of causal inference then combine these two estimates according to specific rules, resulting in predictions that can be compared with behavior in our task.

Below we will briefly describe the important components of our models, and refer interested readers to recent work from Acerbi and colleagues for a much more thorough treatment of this class of models [5]. We begin by describing the cases for location estimation under given causal assumptions (one or two cases), and then describe how these estimates are combined according to different causal inference strategies to produce both judgements about number of targets (unity judgement task) and location of stimulus source(s) (localization task). It is important to note that, while each ‘task’ can be fit independently by different models, the subjects themselves performed the tasks jointly and simultaneously.

#### Fused and Segregated Sensory Localization

For all stimuli, internal representations *x_A_*,*x_V_* are assumed to be corrupted by Gaussian noise, such that 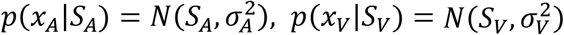 with the *S* term denoting the actual location of the source of the respective stimulus and the *σ* term reflecting the modality specific sensory standard deviation (a free parameter). Estimates about stimulus locations for a given causal structure (c=1, common cause, eq 1; c=2, independent causes, eq 2) and internal representation can be computed via Bayes rule:

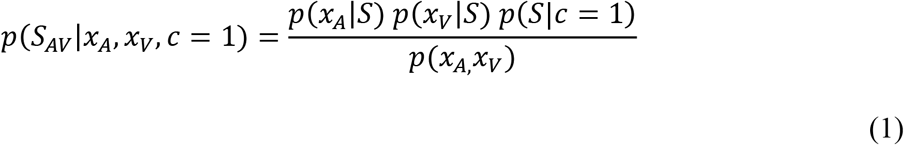

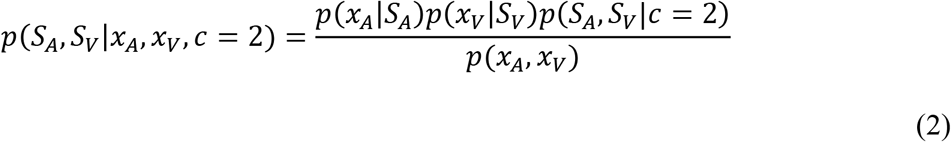

where for the c=1 case the source *S* is assumed to be the same for both the auditory and visual stimuli.

#### Location prior

The subject is assumed to have some prior over possible stimulus locations. A common choice in this type of model is to assume that the subjects have an independent, identical prior over both sensory stimuli,

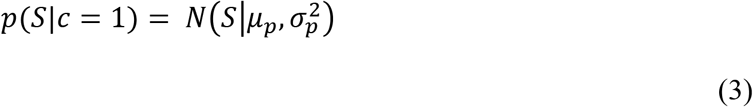

which for the two cause case becomes,

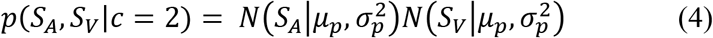

where *μ_p_* is the mean of the prior (here taken as 0) and *σ_p_* is the prior standard deviation. This prior induces a compressive bias which is compatible with many psychophysical results [30].

#### Unity Judgement

The choice of causal inference strategy determines how the observer model decides between the c=1 and c=2 cases when presented with sensory stimuli. In general, this choice can follow either Bayesian principles, non-Bayesian heuristics (i.e. fixed criterion), or strategies which do not actually implement causal inference at all (i.e. forced fusion). There is considerable behavioral work exploring the relative merits of both Bayesian and heuristic forms of causal inference in humans, which is outside the scope of this paper [3–5,48]. Instead, we present a Bayesian form of causal inference and contrast this with a maximally flexible null model which does not perform causal inference at all.

The Bayesian causal inference strategy will compute the posterior probability of the c=1 and c=2 cause cases, given sensory information, as follows,

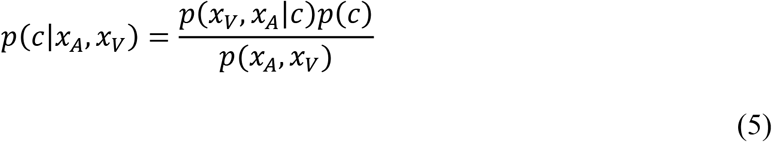

Where p(c) reflects the prior probability of a common cause *p*(*c* = 1) = 1 – *p*(*c* = 2) = *p_common_*, which is left as a free parameter. Because there are only two possibilities for causal state in this paradigm, this can be written as

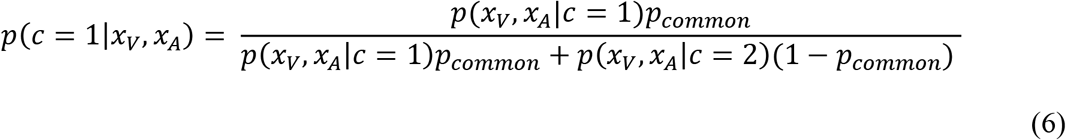

The sensory likelihoods depend on the choice of prior in the previous section, according to

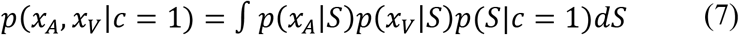

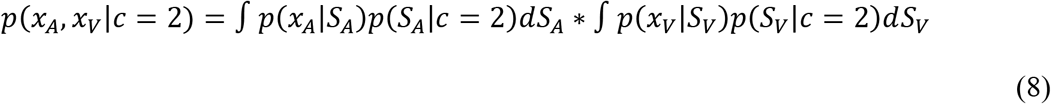

For the simple normal prior these can be solved analytically, but for other forms of prior numerical integration is required. To ensure fairness during the model comparison steps, all likelihoods are computed using the same method (numerical integration).

In order to compare this probability distribution with the binary response distributions (1 or 2 saccades), we must specify a decision rule. For the Bayesian model, we assume subjects report whichever causal scenario has the highest posterior probability. That is,

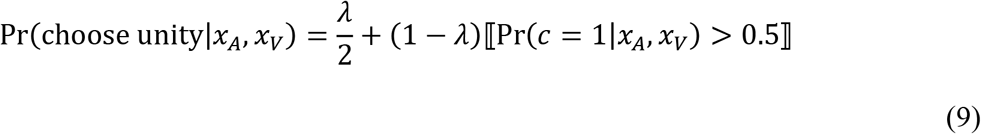

where *λ* represents the lapse rate (the subject randomly makes a response), and 〚·〛 is the Iverson bracket which is 1 when the statement inside is true and 0 otherwise.

The null model for the unity judgement task should allow for automatic fusion, segregation, or some probabilistic choice between the two (potentially reflecting that the subject has learned that a mix of behaviors is required). We implement this by fitting a fixed rate of single saccade responses, equivalent to p_common_, that is common across all combinations of x_A_, x_V_.

#### Localization with Causal Inference

For the localization component of the task, subjects must arbitrate between the two potential models of source location conditioned on number of causes (eqs. 3 and 4). This amounts to reweighting the two estimates according to some weight function that is dependent on the sensory percept,

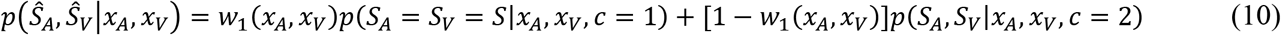

where *w*_1_ defines the decision weight applied to the c=1 condition.

Typically, Bayesian models of causal inference use a model averaging strategy to set these weights. This refers to reweighting the two possibilities according to the posterior probability, such that *w*_1_(*x_A_*,*x_V_*) = Pr(*c* = 1 *x_A_,x_V_*) [2,5,23]. Alternatively, subjects could adopt a model selection strategy. This means they determine which causal structure has the highest posterior probability (eq 6) and then adopt that strategy for the localization component. In this case the weight is equivalent to eq. 9, such that the weight applied to the c=1 condition is 1 when that is the most likely causal scenario and 0 otherwise.

#### Comparing with behavioral data

The above response estimates are dependent on internal variables, x_A_ and x_V_, which are not accessible to the experimenter. To obtain distributions that can be compared with data, eq. 10 must be marginalized across the internal variables:

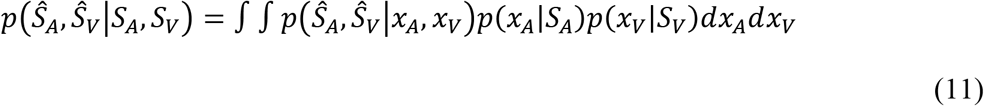

We compute this distribution using numerical integration for each of the 20 combinations of visual and auditory targets, and then use the resulting distributions to calculate likelihood for the purposes of parameter fitting.

For models that assume some form of causal inference, the unity judgement lapse rate, *λ*, will also affect the reported locations (because the subject may make only a single saccade, even though the targets are well separated). To account for this, we included a chance for a single saccade to either the visual or auditory locations rather than the fused location. It was assumed to be equally likely to make an auditory or visual saccade, and the total probability of such saccades was 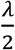.

### Model fitting

Models were fit using a maximum likelihood approach to determine the set of parameters which best explains the provided data. This was accomplished using Matlab’s fminsearch function to minimize the negative log likelihood of the data under each of the above models. The search was initialized using the best starting parameters from a uniform grid search across 10,000 initial parameter settings conducted prior to fitting.

### Model Comparison

We used a Bayesian random-effects model comparison to determine which of the above described models provided the best fits for the observed behavioral data [25]. We performed this model comparison based on the Bayesian Information Criterion (BIC): BIC = −2LL + k x ln(n) where LL is the log-likelihood of the data under the model, k is the number of free parameters, and n is the number of data points. We then determined each models posterior frequency and protected exceedance probability using the variational Bayesian analysis toolbox [26].

## Acknowledgements

We thank Shawn Willett for helpful feedback during the preparation of this manuscript and Jeff Beck for useful discussion about various model features. This work was supported by an NDSEG fellowship to JTM and NIH R01 DC016363 to JMG.

## Author Contributions

**Conceptualization:** Jeff Mohl, John M. Pearson, Jennifer M. Groh

**Data Curation:** Jeff Mohl

**Formal Analysis:** Jeff Mohl

**Funding Acquisition:** Jeff Mohl, Jennifer M. Groh

**Investigation:** Jeff Mohl

**Methodology:** Jeff Mohl, John M. Pearson

**Software:** Jeff Mohl

**Supervision:** John M. Pearson, Jennifer M. Groh

**Validation:** Jeff Mohl

**Visualization:** Jeff Mohl

**Writing – original draft:** Jeff Mohl

**Writing – review & editing:** Jeff Mohl, John M. Pearson, Jennifer M. Groh

